# CytoSimplex: Visualizing Single-cell Fates and Transitions on a Simplex

**DOI:** 10.1101/2023.12.07.570655

**Authors:** Jialin Liu, Yichen Wang, Chen Li, Yichen Gu, Noriaki Ono, Joshua D. Welch

## Abstract

**Summary:** Cells differentiate to their final fates along unique trajectories, often involving multi-potent progenitors that can produce multiple terminally differentiated cell types. Recent developments in single-cell transcriptomic and epigenomic measurement provide tremendous opportunities for mapping these trajectories. The visualization of single-cell data often relies on dimension reduction methods such as UMAP to simplify high-dimensional single-cell data down into an understandable two-dimensional (2D) form. However, these visualization methods can be misleading and often do not effectively represent the direction of cell differentiation. To address these limitations, we developed a new approach that places each cell from a single-cell dataset within a simplex whose vertices correspond to terminally differentiated cell types. Our approach can quantify and visualize current cell fate commitment and future cell potential. We developed CytoSimplex, a standalone open-source package implemented in R and Python that provides simple and intuitive visualizations of cell differentiation in 2D ternary and three-dimensional (3D) quaternary plots. We believe that CytoSimplex can help researchers gain a better understanding of cell type transitions in specific tissues and characterize developmental processes.

**Availability and implementation:** The R version of CytoSimplex is available on Github at https://github.com/welch-lab/CytoSimplex. The Python version of CytoSimplex is available on Github at https://github.com/welch-lab/pyCytoSimplex.

## Introduction

The key goal of CytoSimplex is to quantify and visualize the current state and future differentiation potential of cells undergoing fate transition. Before cells reach their final fates, they often pass through intermediate multipotent states where they have characteristics and potential to generate multiple lineages. Single-cell RNA (scRNA) and epigenome sequencing methods provide temporal snapshots of cells at different moments during cell differentiation. However, since each cell is seen only once and cannot be followed longitudinally, computational approaches must be used to infer where each cell is on its journey. Pseudotime inference and RNA velocity methods have both been used extensively for this task^1–9^. RNA velocity in particular has gained popularity recently. However, the results of these inference methods are often visualized using two dimensional projections of the data derived from principal components (PC), t-distributed Stochastic Neighbor Embeddings (t-SNE) and Uniform Manifold Approximation and Projection (UMAP). While these can be useful visualization tools, they inevitably distort the distance relationships among cells when reducing high-dimensional data to two dimensions^10^. The problem is particularly acute when displaying pseudotime or RNA velocity results; the 2D coordinates are not optimized to show the inferred developmental relationships^11^ and can thus lead to incorrect conclusions about the fate similarity and future potential of multipotent cells. On the other hand, previous computational tools including archetype analysis^12–14^ allow the comparison of similarities between cells and archetypal cells based on a similar concept of simplex. However, these methods do not incorporate predicted future states of cells. Moreover, they can only be applied to scRNA-seq data but not other protocols including single-nucleus ATAC (snATAC) and multi-omic data.

To address these challenges, we developed a new visualization method called CytoSimplex. The key idea of CytoSimplex is to choose clearly defined, interpretable landmarks and plot the positions of transitional cells relative to them. Specifically, we use the molecular signatures of final fates as the vertices of a simplex, then calculate the position of each cell within the simplex. This makes it easier to avoid misleading interpretations due to unclear relationships among cell types on UMAP plots.

## Methods and Results

In CytoSimplex, we model the space of lineage differentiation as a simplex with vertices representing potential terminal fates. A simplex extends the idea of a triangle into any dimension; with a point is a 0D simplex, a line segment is a 1D simplex, a triangle is a 2D simplex, and a tetrahedron is a 3D simplex. A simplex often represents variables that add up to 1 or another fixed value. Intuitively, this is an excellent model for representing cell fate commitment, because there is a small number of cell fates that a given progenitor can produce. This constant sum means the variables cannot change independently, resulting in K-1 degrees of freedom for a K-dimensional simplex. Therefore, a 3D simplex, having two degrees of freedom, can be charted as a 2D triangle. Previously, simplex plots have previously been used in fields like chemistry, geology, and other physical sciences.

We represent the molecular state of a cell as a point located within the simplex. The coordinates of each point (cell) represent its affinity towards each vertex, with these affinities summing to unity. In addition, we incorporate arrows to denote the RNA velocity of each cell towards each terminal state. These simplexes and arrows together serve as cell-wise representations of putative transcriptomic and epigenomic transitions towards a specific trajectory, guided by the selection of putative terminal fates. CytoSimplex takes single-cell RNA and/or single-cell ATAC count matrices, a cell-cell transition graph (such as a result from RNA velocity analysis^1,2^), and cell type annotations as input (**Fig. 1A, left**). To generate simplex dot plots, for each dataset and chosen terminal cell type, we derive a distance metric by selecting the top 30 differentially expressed marker genes by Wilcoxon test, calculating the Euclidean distance from each cell to the mean-centroid of each cluster, and normalizing the distances to a sum of unity as the barycentric coordinates. We then project them onto a Cartesian coordinate system to generate the simplex plot. For visualizing dynamic velocity vectors, we first derive a cell-cell transition graph by performing RNA velocity methods including MultiVelo^3^, VeloVAE^4^, and scVelo^1^. We then calculated the future differentiation potential of each cell by taking the average of the edge weights between each cell and the cells in each terminal cluster. For a more concise view, we binned cells into square or cube grids, aggregated the weights by the total number of cells in each grid, and normalized them to a sum of unity. Finally, the velocity was depicted as arrows pointing from the centroid of each grid to each vertex, with arrow length representing the normalized weights (**Fig. 1A, right**).

**Figure 1.**
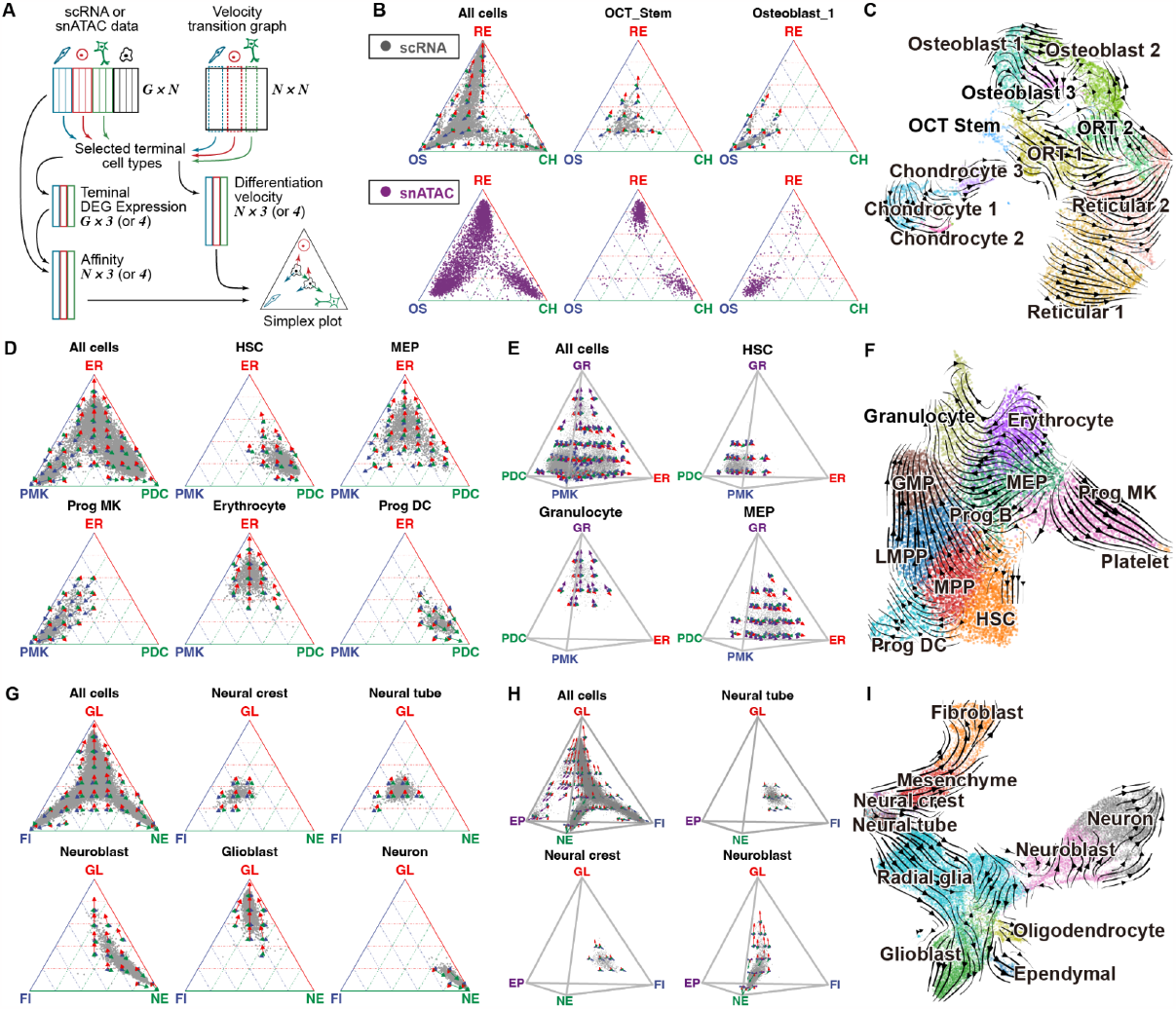
CytoSimplex shows multi-lineage differentiation potential of progenitor cells in different organs. **(A)** Input and output of CytoSimplex. CytoSimplex takes a gene-by-cell RNA and/or ATAC count matrix, RNA velocity transition matrix, and cluster assignments as input, and generates ternary or quaternary plots. The coordinates of each cell within the simplex depict its affinity towards each vertex, with dynamic arrows inferred from RNA velocity analysis indicating the cell’s differentiation potential towards each terminal fate. The color coding of both the arrows and vertices corresponds to the final fates selected. Together, these simplexes and arrows provide an intuitive visualization of cellular trajectories. **(B)** Ternary plots of representative cell types in embryonic mouse bone marrow generated from scRNA and snATAC data. OCT stem cells demonstrate transcriptomic potential towards all three terminal fates, while also showing epigenomic potential towards Reticular and chondrocyte cells. Red arrow and axis: Reticular cluster (RE). Blue: Osteoblast cluster (OS). Green: Chondrocyte cluster (CH). **(C)** RNA velocity of major cell types in mouse bone marrow. Corresponding to panel (A) and (B), RNA dynamic vectors originate from OCT-stem cells and point to other differentiated cell types, suggesting OCT-stem cells as the cell-of-origin cluster. **(D, E)** Ternary and quaternary plots of human hematopoietic stem and progenitor cells (HSPC). Corresponding to their locations on the UMAP plots in (E), progenitor cell types including HSC and MEP show distinct transcriptomic similarities to all terminal fates, while showing identical differentiation potential towards all vertices, regardless of in the ternary or quaternary plot. HSC: Human hematopoietic stem, MEP:Megakaryocyte-erythrocyte progenitor. Red arrow and axis: Erythrocyte cluster (ER). Blue: Progenitor Megakaryocyte cluster (PMK). Green: Progenitor Dendritic cluster (PDC). Purple: Granulocyte cluster (GR). **(F)** RNA velocity of major cell types in HSPC data. HSC and MEP, functioning as progenitor cell types, serve as two distinct cell-of-origin clusters. **(G, H)** Ternary and quaternary plots of major cell types in mouse brain atlas. Neural tube and neural crest cell, as two topologically close stem cell types, share similar transcriptomic similarities and differentiation potential towards all fates in ternary and quaternary plots. GL: Glioblasts, FI: Fibroblasts, NE: Neurons, EP: Ependymal cells. **(I)** RNA velocity of major cell types in mouse brain atlas. Neural tube and neural crest cell serve as root cells of distinct differentiation trajectories while also sharing a similar location in UMAP plot.

CytoSimplex is different from prior methods due to its novel approach for showing both the current and future state of each cell. Previous methods like archetype analysis^12–14^—which resembles the simplex concept—allow the comparison of similarities between cells and terminal cell types in a low-dimensional space defined by archetypal cells, and therefore helps determine if cells are multitasking during development. Yet CytoSimplex sets itself apart because it not only shows the relationships among current cell states, but also the fate potential and predicted future states of the cells. This unique feature allows CytoSimplex to accurately represent cell lineage transitions dynamically and intuitively, measuring how much potential each cell has of differentiating into each terminal fate.

To enhance user-friendliness, we have developed an open-source package in both R and Python. This package is well-structured and specially tailored for researchers who are not familiar with programming. On top of that, CytoSimplex is versatile, compatible with multiple sequencing protocols. Our method takes any cell-by-feature matrices accompanied by cell-cell transition graphs and cell type annotations as input. We aim to contribute to the bioinformatics community by distributing our package to the widest range of researchers.

To illustrate how CytoSimplex can be used, we first employed our method on mouse bone marrow stromal cells (BMSCs) scRNA-seq and snATAC-seq data from our recent papers^15–18^. This BMSC population encompassed large groups of osteoblastic cells (Osteoblasts, OS), pre-adipocyte-like reticular cells (Reticular cells, RE), separate clusters of chondrocytes (Chondrocytes, CH), and a putative stem cell cluster with osteoblast-chondrocyte transitional identities (OCT stem cells) (**Fig. 1C**). Velocity analysis suggests that OCT stem cells provided a robust cellular origin of osteoblasts, reticular cells, and their intermediate-state cells. CytoSimplex confirms but deepens the story by demonstrating the transcriptomic similarity and differentiation potential of each cell in the dataset. Specifically, in our simplex plots, the majority of OCT stem cells stay in the middle of the simplex and have almost equal similarity to each fate (**Fig. 1B, OCT stem**), while also demonstrating “trilineage” potential toward all three fates indicated by arrows pointing towards three vertices (**Fig. 1B, OCT stem**). In contrast, the chromatin accessibility profiles of OCT stem cells reveal “trilineage” potential that aligns with the transcriptomic profiles, but with a greater bias toward the reticular and chondrocyte fate (**Fig. 1B, snATAC**). As comparison, cells within each terminal cluster exhibited strong RNA and ATAC similarities to their respective fates (**Fig. 1B, Supplementary Fig. S2 and S3**). Therefore, CytoSimplex can not only detect the multipotency of a potential stem cell population, but also uniquely unveiled the disagreements between the transcriptomic and epigenomic developmental direction of OCT-stem cells.

Subsequently, we applied our method to human hematopoietic stem and progenitor cells (HSPCs) single-nucleus RNA+ATAC multiome data employed in our published study^3^. This dataset comprises two distinct global populations of stem cells, namely hematopoietic stem cells (HSCs) and multipotent progenitors (MPPs). These cells are capable of differentiating into three main lineages: myeloid lineage, lymphoid lineage, and dendritic lineage. Each lineage consists of various progenitor cell types including megakaryocyte-erythroid progenitors (MEP), as well as more differentiated blood cells such as granulocytes (GR), erythrocytes (ER), and platelets. Analysis of the RNA velocity streamline plot based on UMAP reveals a close proximity between HSCs and progenitor dendritic cells (PDCs) on the developmental trajectory, while MEPs exhibit similarities to both ER cells and the combined population of platelets and progenitor megakaryocytes (PMKs) (**Fig. 1F**). Consistently, in terms of transcriptomic similarity, our ternary plots confirm these findings, with HSCs staying close to the vertex of PDCs, while MEP cells predominantly align with the ER and PMK fates. However, CytoSimplex also highlights divergence between HSCs and MEP cells. HSCs exhibit high differentiation potential across all vertices, while MEP cells tend to favor the ER and PMK vertices, which belong to the same myeloid lineage (**Fig. 1D**). Furthermore, the quaternary plots unveiled the same story when we included granulocytes (GRs), which are differentiated from granulocyte-monocyte progenitors (GMPs) instead of MEPs, as an additional vertex (**Fig. 1E, Supplementary Fig. S1A**). We also found that except the ones serving as the vertices, none of the progenitor cell types reach to the endpoints of any vertices in both ternary and quaternary plots (**Supplementary Fig. S4 and S5**). This aligns with our biological expectation that the majority of differentiating progenitor cells are multipotent cells that have not yet reached their final fates. Therefore, our simplex method provides a clearer understanding of the intricate relationship between progenitor cells from multiple lineages and also highlights their pluripotent state.

Moreover, we applied our method to part of a mouse brain atlas. Our previous work performed velocity analysis and demonstrated three distinct differentiation paths including glioblasts, fibroblasts, and neurons^4,19^. The progenitor cells, namely neural crest and neural tube cells, can develop into multiple descendants. In particular, neural crest cells develop into fibroblast cells via mesenchymal cells, while neural tube cells end up becoming both neuronal and non-neuronal cells, including oligodendrocytes, ependymal and glioblast cells. While the velocity stream generally follows the direction of these differentiation paths, some streamlines can be misleading due to distorted cell-cell relationships introduced by dimension reduction^10^. For example, UMAP velocity of neuroblast cells deviates significantly from major paths from radial glia cells and towards neurons (**Fig. 1I**). In contrast, our ternary plot indicates an evident overall trend of neuroblast cells differentiating into the neuron fate, with some cells showing similarities towards the glioblast fate (**Fig. 1G**). We further highlight that CytoSimplex captures high similarity in differentiation tendency between two topologically close stem cell types, the neural tube and neural crest cell, while still capturing the subtle distinctions. Specifically, neural crest cells have a higher tendency to undergo fibroblast cell transformation in comparison to neural tube cells. To demonstrate a four-lineage viewpoint, we introduced Ependymal as an additional end fate (**Fig. 1H, Supplementary Fig. S1B**). Progenitor cells, such as neural crest and neural tube cells, exhibit a similar neutral affinity towards all four vertices. CytoSimplex also validates clear differentiation trends indicated by RNA velocity streams, such as mesenchymal cells predominantly differentiated towards the fibroblast fate (**Supplementary Fig. S6**). Additionally, cells in each terminal cluster demonstrated the strongest similarities and differentiation trend towards their own fates.

In conclusion, CytoSimplex is an accessible and innovative tool that provides simple and intuitive visualization of complex cell transition trajectories. Its unique ternary or quaternary plots provide cell-wise information by combining cellular transcriptome or epigenomic signatures with RNA velocity-inferred differentiation potential, which allows researchers to achieve a significantly higher resolution analysis compared to commonly used visualization methods. By utilizing CytoSimplex, researchers can easily focus on each lineage within a complex dataset and discern minor heterogeneities between intriguing cell populations, such as stem cells and progenitor cells. Furthermore, CytoSimplex’s implementation in R and python enables researchers to streamline their analysis with other single-cell analysis tools, improving their ability to analyze complicated datasets. We believe that CytoSimplex can help advance the understanding of biological processes underlying cell differentiation.

## Supplementary Figures

**Supplementary Figure 2.**
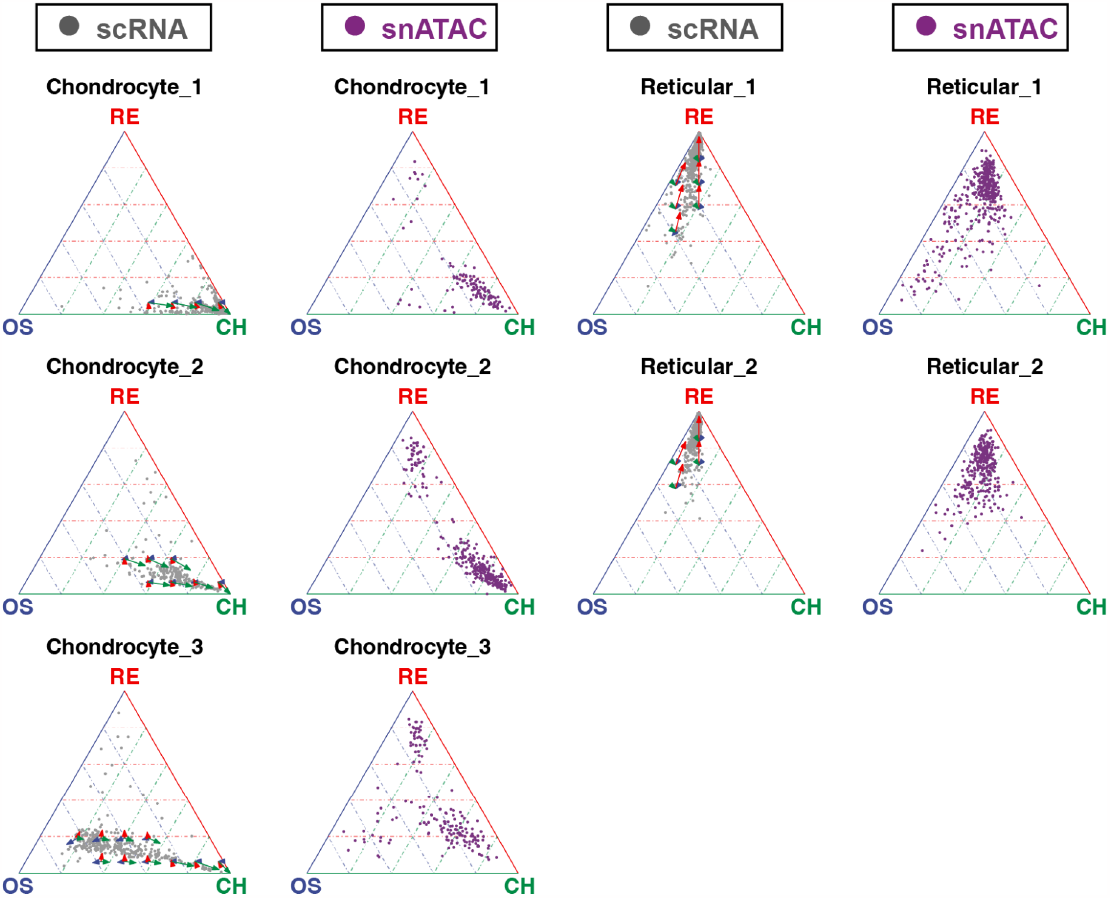
Ternary simplex and velocity analyses of BMMC data. In the first and third columns, cells colored in gray demonstrate their transcriptomic affinities towards three selected vertices, whereas the arrows show the future differentiation potential of the cells. In the second and fourth columns, cells colored in purple demonstrate their epigenomic affinities towards three vertices. Red arrow and axis: Reticular cluster (RE). Blue: Osteoblast cluster (OS). Green: Chondrocyte cluster (CH).

**Supplementary Figure 3.**
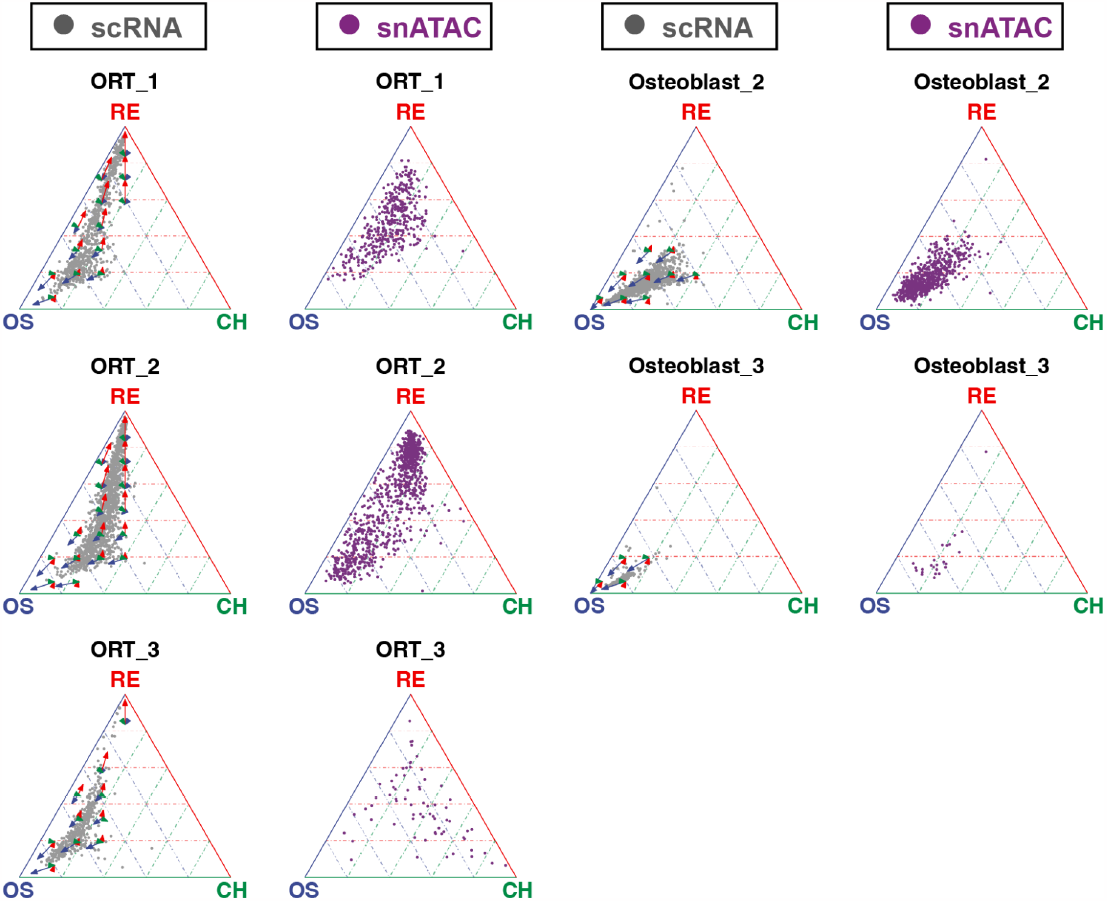
Ternary simplex and velocity analyses of BMMC data. Continuing from Supplementary Figure 1, this figure includes the remaining clusters from BMMC data. Red arrow and axis: Reticular cluster (RE). Blue: Osteoblast cluster (OS). Green: Chondrocyte cluster (CH).

**Supplementary Figure 4.**
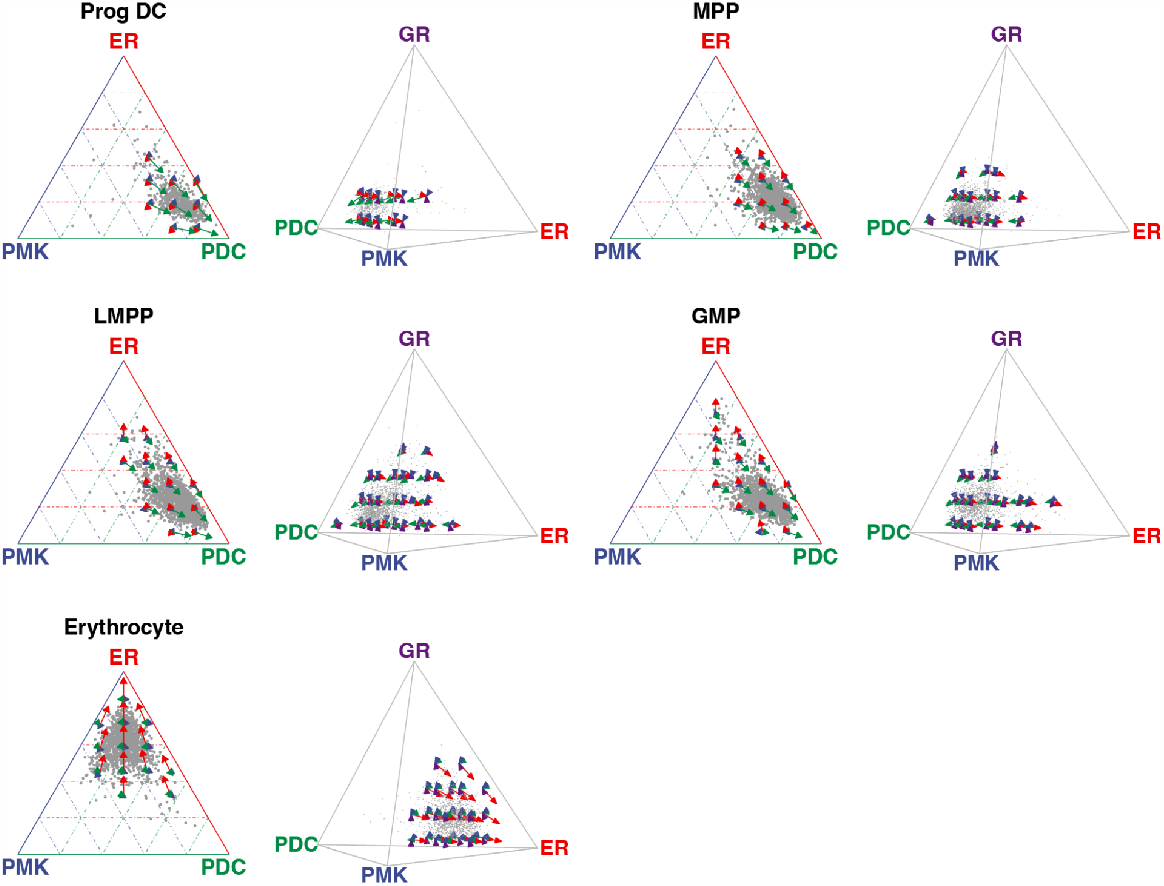
Simplex and velocity analyses of HSPC data. Cells colored in gray demonstrate their transcriptomic affinities towards three selected vertices, whereas the arrows show the future differentiation potential of the cells. Ternary simplex plots are shown in the first and third columns, while quaternary simplex plots are shown in the second and fourth columns. Red arrow and axis: Erythrocyte cluster (ER). Blue: Progenitor Megakaryocyte cluster (PMK). Green: Progenitor Dendritic cluster (PDC). Purple: Granulocyte cluster (GR).

**Supplementary Figure 5.**
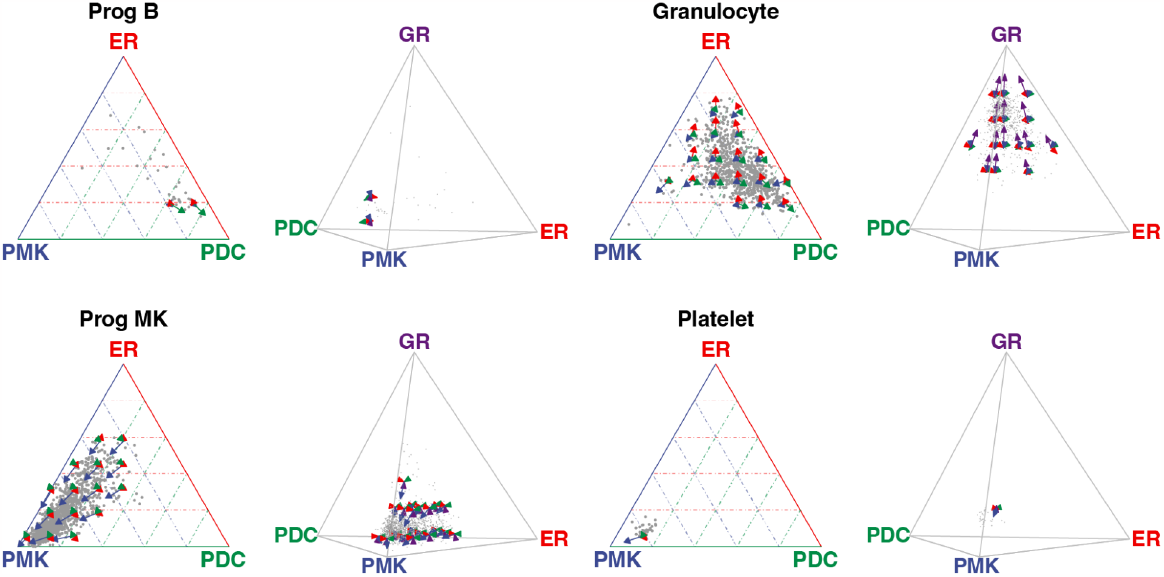
Simplex and velocity analyses of HSPC data. Continuing from Supplementary Figure 3, this figure includes the remaining clusters from HSPC data. Red arrow and axis: Erythrocyte cluster (ER). Blue: Progenitor Megakaryocyte cluster (PMK). Green: Progenitor Dendritic cluster (PDC). Purple: Granulocyte cluster (GR).

**Supplementary Figure 6.**
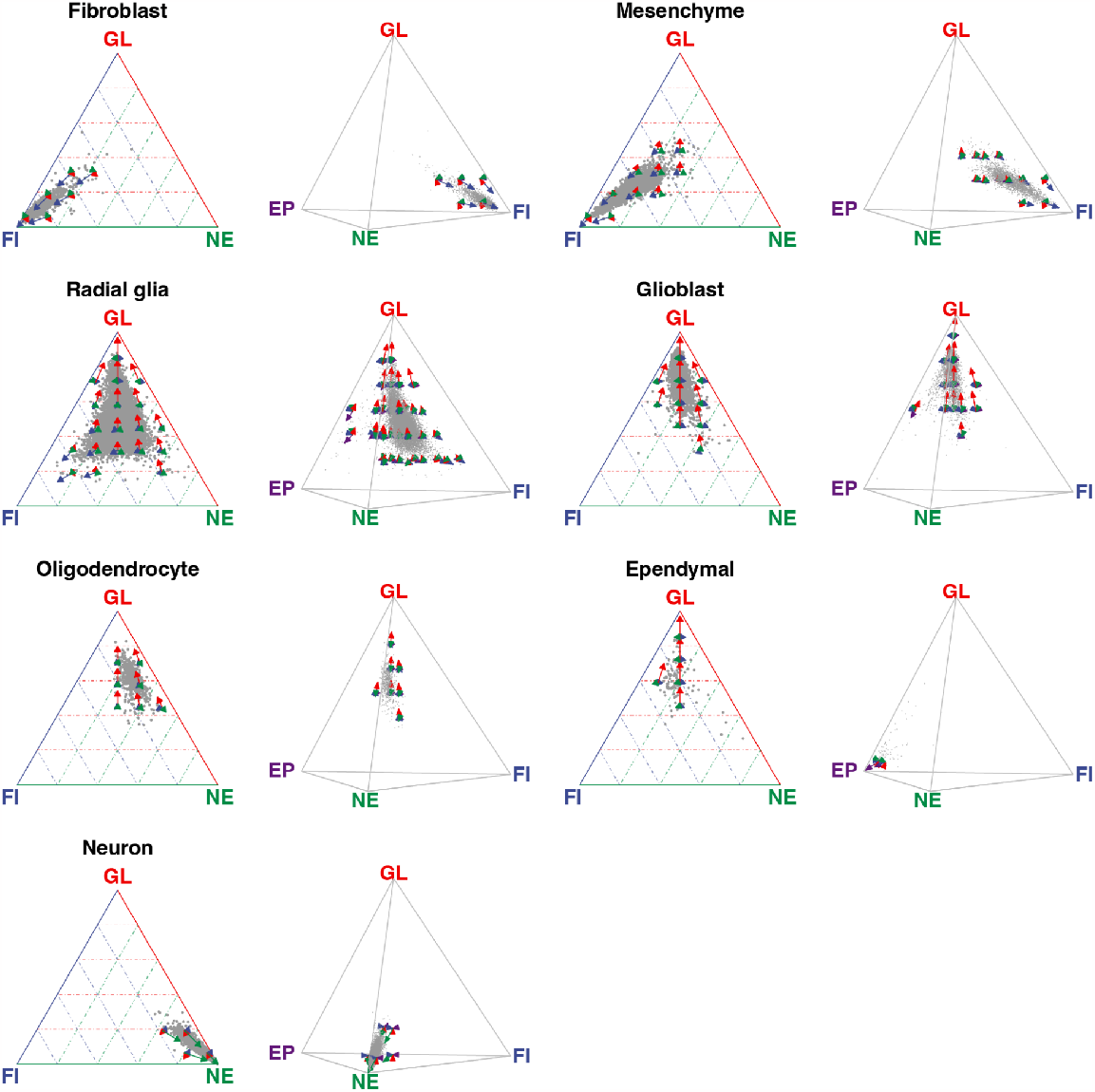
Simplex and velocity analyses of mouse brain atlas data. Cells colored in gray demonstrate their transcriptomic affinities towards three selected vertices, whereas the arrows show the future differentiation potential of the cells. Ternary simplex plots are shown in the first and third columns, while quaternary simplex plots are shown in the second and fourth columns. Red arrow and axis: Glioblast cluster (GL). Green: Neuron cluster (NE). Blue: Fibroblast cluster (FI). Purple: Ependymal cluster (EP).

