## Supplementary Fig. S1 for "CytoSimplex: Visualizing Single-cell Fates and Transitions on a Simplex"

**Supplementary Figures**

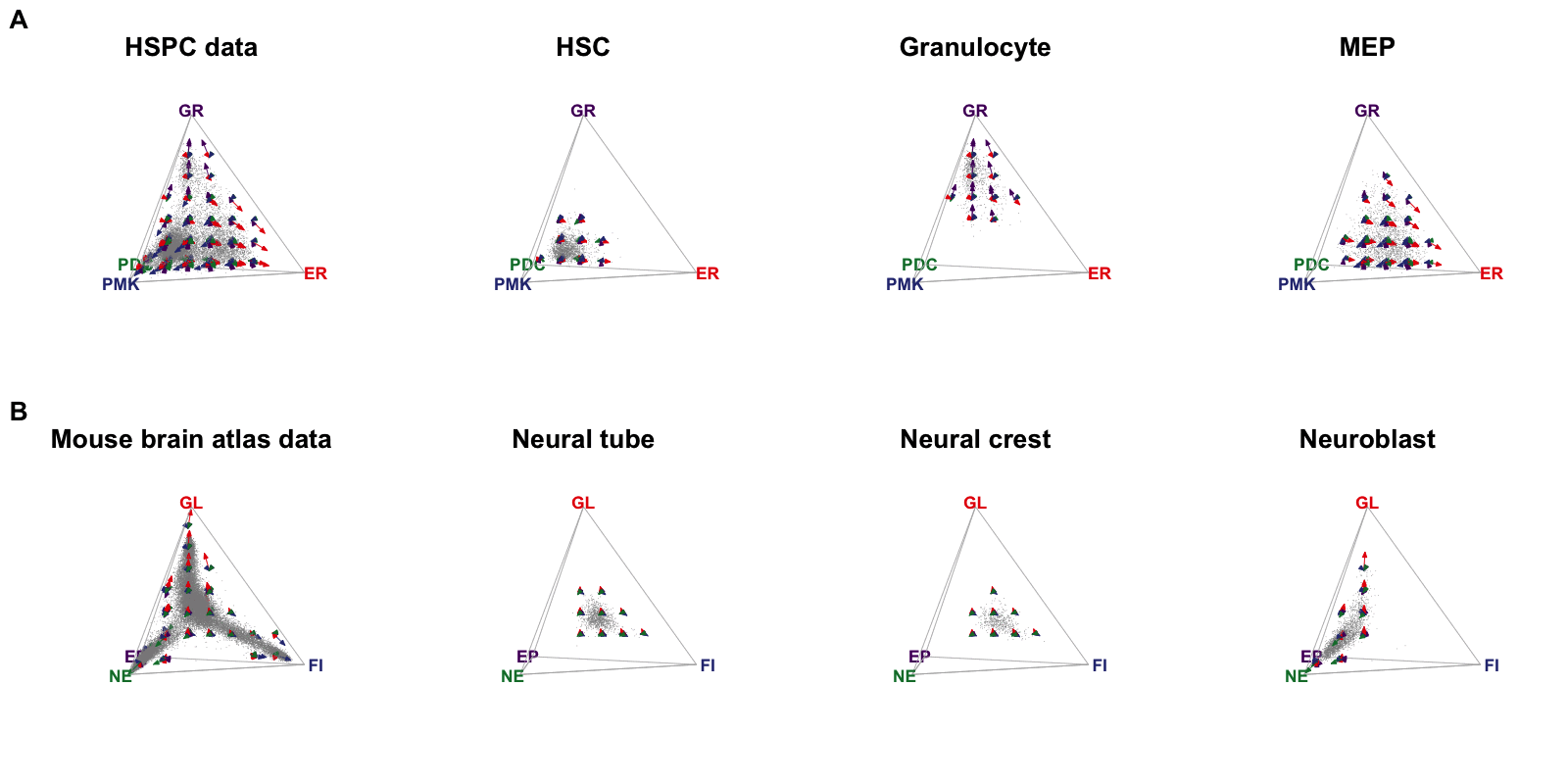

**Supplementary Figure 1.** Animated 3D quaternary simplex plots of HSPC and mouse brain atlas data. Continuing from Figure 1, these animated quaternary simplex plots of HSPC **(A)** and mouse brain **(B)** data provide a full 360-degree view and therefore offer a more detailed demonstration of the relationship between selected cell types and terminal cell fates. For HSPC data, Red arrow and axis: Erythrocyte cluster (ER). Blue: Progenitor Megakaryocyte cluster (PMK). Green: Progenitor Dendritic cluster (PDC). Purple: Granulocyte cluster (GR). For mouse brain atlas data, Red arrow and axis: Glioblast cluster (GL). Green: Neuron cluster (NE). Blue: Fibroblast cluster (FI). Purple: Ependymal cluster (EP).
